# PlasForest: a homology-based random forest classifier for plasmid detection in genomic datasets

**DOI:** 10.1101/2020.10.05.326819

**Authors:** Léa Pradier, Tazzio Tissot, Anna-Sophie Fiston-Lavier, Stéphanie Bedhomme

## Abstract

Plasmids are mobile genetic elements that often carry accessory genes, and are vectors for horizontal transfer between bacterial genomes. The detection of plasmids in large sets of genomes is crucial to analyze their spread and quantify their role in bacteria adaptation and particularly in antibiotic resistance genes propagation. Several bioinformatics methods have been developed to detect plasmids. However, they suffer from low sensitivity (*i.e*., most plasmids remain undetected) or low precision (*i.e*., these methods identify chromosomes as plasmids), and are overall not adapted to identify plasmids in whole genomes that are not fully assembled (contigs and scaffolds). Here, we present PlasForest, a homology-based random forest classifier identifying bacterial plasmid sequences in unassembled genomes. This tool is based on the determination of homologies against a database of plasmid sequences, which allow a random forest classifier to discriminate plasmid contigs. Without knowing the taxonomical origin of the samples, PlasForest identifies contigs as plasmids or chromosomes with an accuracy of 98%. Notably, it can detect 96% of plasmid contigs over 50kb with 3.3% of false positives. PlasForest outperforms other currently available tools on test datasets by being both sensitive and precise. We implemented this tool in a user-friendly pipeline that can identify plasmids in large datasets in a reasonable amount of time.

## 1. Introduction

Plasmids are extra-chromosomal fragments of DNA that replicate autonomously in the host cell. They often carry genes that can provide a benefit under specific environmental conditions (1). These mobile genetic elements remain a major biological concern for health and agriculture policies due to their ability to accumulate and spread resistance genes. Indeed, the frequency of plasmids, and of the resistance genes they carry, can increase quickly in populations thanks to their high mobility both within hosts (through chromosomal integration) and between hosts (through horizontal gene transfer, hereafter HGT). Due to their high mobility and the function of the genes they carry, plasmids have a great ecological importance in many bacterial communities.

Until recently, the identification and isolation of these mobile genetic elements was limited to a narrow subsample of the bacteria diversity. Most past studies have only focused on specific species-plasmid associations of medical or agronomic interest (2, 3). On the one hand, the plasmid characteristics determining their role in resistance propagation were established by phenotypic approaches, which require the focal strain to grow on culture media. These approaches could assess the ability of genetic elements to transfer small conjugative plasmids carrying selectable markers into a new recipient, *i.e*. transfer, replication and expression of selectable markers (4), but were useless for non conjugative plasmids. On the other hand, PCR-based detection methods (5) could detect the number of copies for specific plasmid sequences, but did not allow to understand the ecological characteristics of mobile genetic elements, as for example their host range (5).

The fast development of sequencing technologies and reduction of sequencing costs led to the rapid increase of available genomic and metagenomic data sets. This material contains a vast amount of information on plasmid diversity, plasmid host-range, resistance conferred to specific host taxa, etc. that could allow to better understand the circulation and spread of plasmids and the genes they carry. Accessing this information requires new tools to process raw data such as contigs and scaffolds and metagenomes from environmental samples to identify plasmids sequences.

Well-defined plasmids can be identified through homology search (*e.g*., PlasmidFinder (6), PLACNET, PLACNETw (7)). These programs basically look for similarities between a query sequence and a local database. Query and subject sequences are usually quite long (at least several hundreds of bases), so the probability of finding similarities by chance (and thus to wrongly identify a sequence as plasmid) is very low. Homology search is usually very precise and reliable. However, current data limitations can substantially decrease the sensitivity of this method. Indeed, it requires an exhaustive database composed of a wide variety of genomics data, enough to cover many taxa, and even if this is the case, rare plasmids may not be identified. In particular, the range of taxa in which a plasmid can be replicated and maintained (*i.e*., its host range (5)) can greatly vary: some being restricted to a few close species, and others consisting in a wide range of taxa across the phylogeny. Thus, plasmids with a narrow host range (especially when only present in uncultivated species) could be nearly impossible to identify through homology.

In the absence of an exhaustive database, an accurate identification method requires to define broader associations with plasmids. This can be achieved by the reconstruction of plasmids through homology-based graphs (PLACNET, PLACNETw (7)): the reads which are homologous to the same reference plasmid sequences are likely to be part of the same plasmids. This approach can be very successful if the generated graphs are manually pruned by an expert user but this strongly impedes the possibility to apply this method to large datasets.

Other algorithms identify plasmids in draft genomes and metagenomes without relying on homologies, but rather on k-mers frequencies: PlasFlow (8); cBar (9); PlasmidSeeker (10). Scoring sequences based on their k-mers frequencies is quick, automated, and adaptable to various data, and may result in valuable predictions (*e.g*., more than 95% of correct predictions for PlasFlow (8)). However, the distribution of k-mers cannot be estimated with precision in a short contig. Even when using short k-mers (*e.g*., 7-mers for PlasFlow (8)), precision may be lower for the shortest contigs, and most k-mer-based methods systematically rule out contigs below 1kb.

Overall, the current plasmid identification methods are limited, in either precision or sensitivity. These limitations are not problematic for certain applications: for example k-mers based methods are well adapted to detect plasmid sequences in metagenomics samples containing many uncultivable species and homology-based approaches are well suited if users focus only on a few plasmids of interest. Yet, no tool is currently available to classify, with both high precision and sensitivity, a large data set of sequences as plasmid or chromosome. Such a tool would be very useful for example to monitor the spread of a gene family (*e.g*., antibiotic resistance genes) between genomes. Indeed, 90% of available genomes on public repositories (up to 100% for species with few sequenced genomes) are unassembled genomes (contigs or scaffolds). It is thus not possible to rely on the size of the genomic elements and their content in specific sequences (as ribosomal RNA genes or origin of replications) to classify them as chromosome or plasmids and an accurate plasmid identification method is required.

Here, we present PlasForest, a new tool for classifying contigs in genomic datasets as plasmid or chromosome DNA. Contrary to classical homology search, our method does not attempt to assign query contigs to specific subject sequences, but rather to sort contigs in plasmid and chromosome sequences through a machine learning algorithm fed with parameters extracted from an homology search against an exhaustive plasmid database, as well as other variables (*e.g*., contig size and %G+C content). PlasForest can discriminate plasmids from chromosome sequences with an overall accuracy of 98% for any bacterial contig in genomic datasets. In particular, PlasForest is able to detect up to 78% of plasmid contigs under 5kb with less than 11% false-positives and up to 96% of plasmid contigs over 50kb with less than 0.5% false-positives. Compared to other currently available tools, PlasForest has a significantly better capacity to correctly identify plasmids from chromosomes. We implemented this tool in a user-friendly pipeline able to identify plasmids in large datasets in a reasonable amount of time.

## 2. Material and methods

The aim of PlasForest is to combine both the high precision of homology search with the broad sensitivity of signature-based classifiers in order to discriminate contigs of plasmid origins from contigs of chromosomes. We trained a classifier for which decision relies on the homology of sequences with a large database of plasmid sequences. We simulated contigs by randomly cutting assembled genomes, to construct both a dataset to train the classification algorithm (the training set) and a dataset to measure the classification performance (the testing set). We then compared the classification performance to other plasmid identification tools.

### 2.1 Data collection

#### 2.1.1 Plasmid database

All bacterial plasmid sequences were downloaded from the NCBI RefSeq Genomes FTP server (ftp://ftp.ncbi.nlm.nih.gov/genomes/refseq; September 1st, 2019). This database is composed of 36,450 sequences that we used as reference for homology seeking *via* BLAST tool (11) (e-value < 10^−3^).

#### 2.1.2 Training, testing, and external validation datasets

To train the classifier and measure its performance, we randomly sampled 14,909 bacterial genomes classified as ‘complete’, downloaded from the NCBI Refseq Genomes FTP server (ftp://ftp.ncbi.nlm.nih.gov/genomes/refseq). This set of genomes was then split in a *training set* (about 46% of the sequences), a *testing set* (about 23% of the sequences) and an *external validation set* (about 33% of the sequences) (see Fig. 1). These sets were built such that the collections of genera present in the training and testing sets were largely overlapping whereas the collections of sequences of the training and testing sets on one side and the external validation set on the other side had no overlap, based on NCBI taxonomy data (see Fig. 1). The *external validation dataset* was used to validate our classification method.

**Figure 1:**
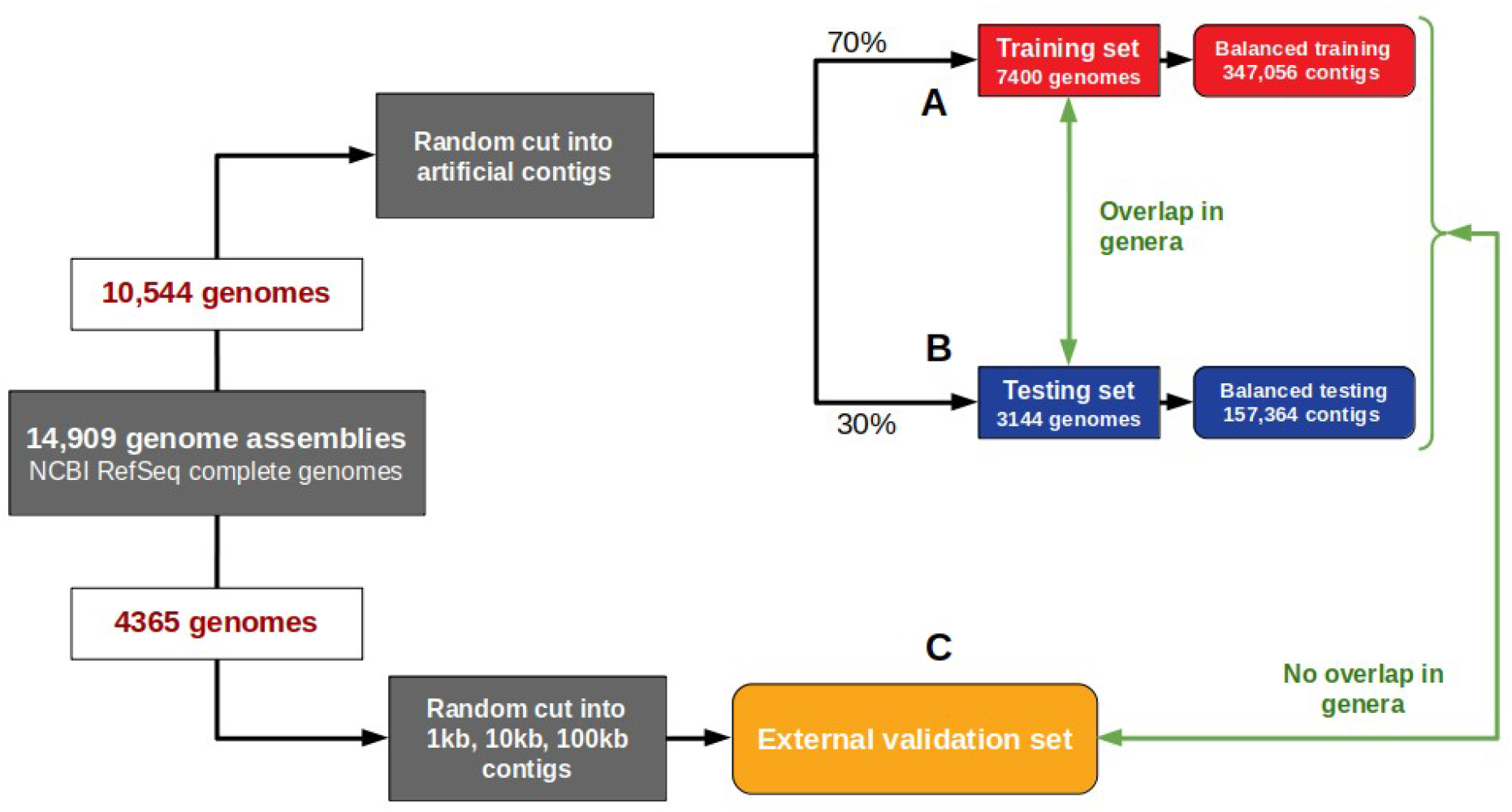
Datasets and application of a hold-out method for supervised learning. Schematic representation of the processes that allow to generate three datasets used to build PlasForest and to benchmark its performances. 14,909 bacterial genomes from NCBI Refseq Genomes FTP server were randomly cut into contigs, and were distributed into the following datasets. (A) *(Balanced) training set* 70% of the initial 10,544 genomes assemblies. This dataset is used to train the random forest classifier. (B) *(Balanced) testing set* contains 30% of the contigs. This dataset is used to benchmark the performances of PlasForest against other existing identification methods. (C) The *external validation dataset* contains 33% of the 14,909 initial genomes. It was used to validate our classification method.

To mimic the sequence material on which PlasForest will be applied (contigs from unassembled whole genomes), the empirical distribution of contig sizes was established from more than 100,000 Refseq unassembled genomes (see Supplementary information, Tab. S1 for the chosen distribution). This distribution was then artificially recreated from complete genomes in the training and testing sets by cutting plasmids and chromosomes at random locations and keeping a defined number of each contig size in plasmids and chromosomes. Only contigs larger than 50bp were kept, since most current sequencing approaches do not produce shorter reads (12). We ended up with approximately 70% of the generated contigs (552,410 contigs coming from 7,400 genomes) to train PlasForest. Genome annotations were used to identify contigs as plasmids or chromosomes. The remaining 30% of the generated contigs were used as a testing set. In these two datasets, plasmid contigs were not at the same frequencies for all the contig sizes (*e.g*., >30% under 1kb and 2% over 100kb). This could have led to an artificial detection bias based on contig size (*e.g*., a better identification of small plasmid contigs). Thus, we split initial datasets into contig size categories (50bp to 1kb, 1 to 2kb, 2 to 5kb, 5 to 10kb, 10 to 50kb, 50 to 100kb, and over 100kb), and randomly removed plasmid or chromosome contigs from each category to keep the fraction of plasmid contigs constant (around 10%) across contig sizes. These new datasets are thereafter called *balanced datasets* (*i.e*., balanced training set and balanced testing set respectively, Supplementary information, Tab. S2A, Tab. S2B).

In order to clearly assess the impact of contig size on the performance of the classification algorithm, the genomes *of the external validation set* were randomly cut into pieces of 1kb, 10kb, 100kb (Supplementary information, Tab. S3). The lists of contigs used to build the balanced training set, the balanced testing set, and the external validation set are available in supplementary material (Tab. S2, Tab. S3).

### 2.2 Construction of PlasForest

#### 2.2.1 Extraction of the features

All contigs were compared against the plasmid database defined in section 2.1.1 using BLASTn (9). Pair-alignments with homologous sequences (hereafter referred to as “hits”) were kept if their e-value was below 10^−3^. For each contig and homologous sequence, we computed *overlap* as the fraction of the query contig aligning to the homologous sequence hit. The G+C content of all contigs was computed with the function *SeqUtils.GC* from the Biopython library in Python 3.6 (13).

Our aim is not to assemble plasmids (or to assign contigs to precise replicons), but to identify contigs that originate from plasmids. This motivates for a distinct design from other homology-based approaches. By combining both homology search and measures of nucleotide composition, we aim to obtain a strong distinction between plasmids and chromosomes. We therefore selected features as follows. (i) ***Maximal overlap*** was measured among hits in the subject database, because we expect that query plasmid contigs should form longer alignments with sequences from the plasmid database than query chromosomes. (ii) ***Contig size*** was taken into account as short contigs align more often than large contigs with the subject database. (iii) The ***number of hits***, the ***average overlap***, the ***median overlap***, and the ***variance of overlaps*** provide other parameters of the distribution of overlaps among hits, that may help distinguish between chromosomes and plasmids. Indeed, due to recombination events, one may expect that query chromosome contigs will align with subject plasmids, but more rarely than query plasmid contigs. (iv) Finally, the ***G+C content*** was also included, as the nucleotide composition of plasmids are most often different from those of chromosomes (14). This set of features used to train the classifier is schematically displayed in Fig. 2A. These features were found to be very different for plasmid contigs and chromosome contigs of the balanced training set (see Fig. 2B, 2C).

**Figure 2:**
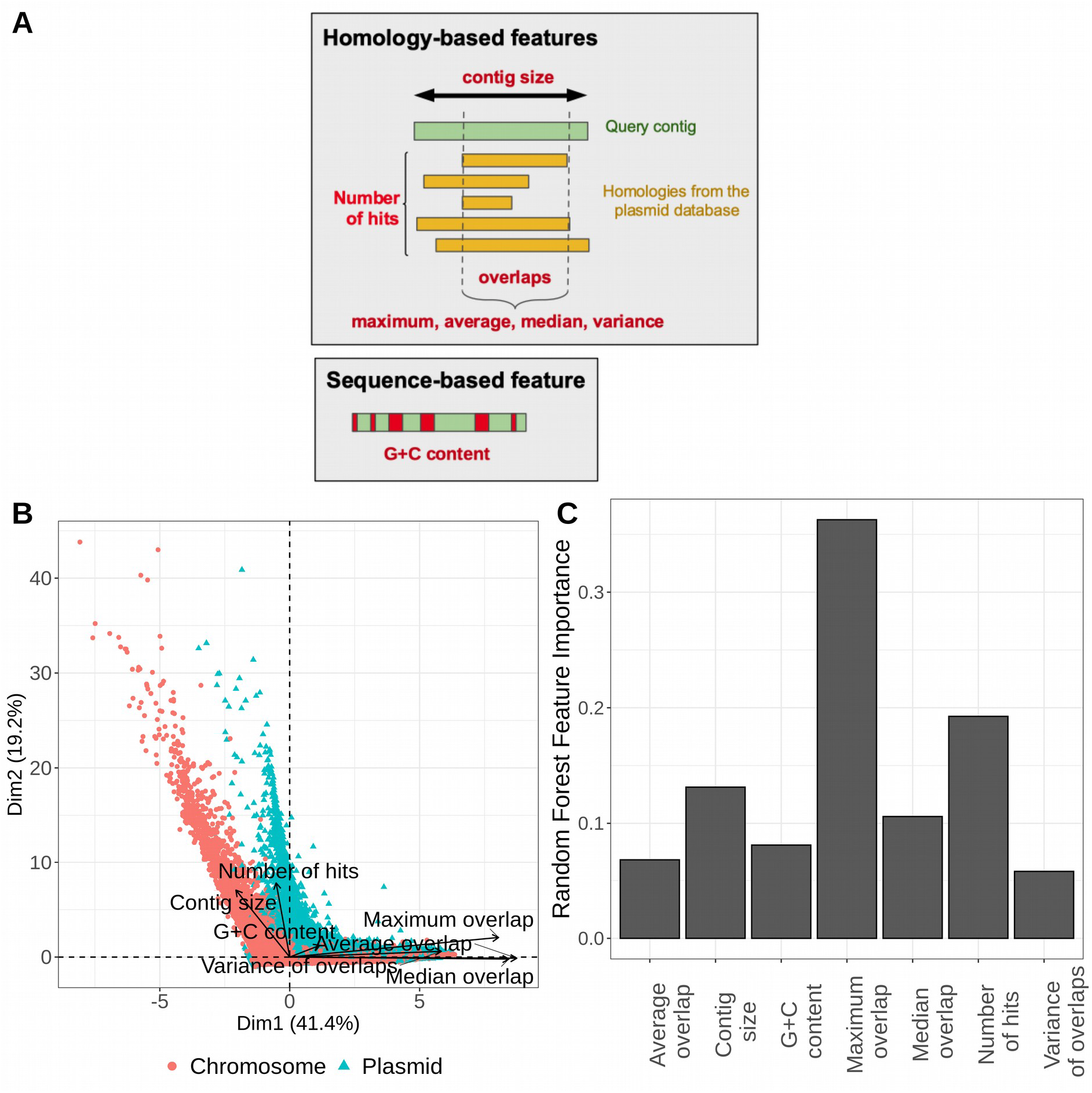
Chosen features and their importances in the classification process. (A) Schematic representation of the features extracted from contigs, including homology-based features (number of hits, maximum overlap, average overlap, median overlap, variance of overlaps, contig size) and sequencebased feature (G+C content). (B) Principal component analysis of the features for contigs in the balanced training set: contigs of chromosomal origin are displayed as red dots, and contigs of plasmid origin as blue triangles. The chosen features allow to draw two different distributions for plasmids and chromosomes in the training set. We use these features to train a classifier. (C) Impurity-based feature importance computed with *scikit-learn* library for the seven features kept in the classifier

#### 2.2.2 Training of the classifier

We decided to extract the differences in the features of plasmid contigs and chromosome contigs through a random forest classifier. This approach relies on a multitude of independent decision trees, which allows for a reduction of individual error (15). The aim was therefore to build a model able to predict, from the extracted features, whether a contig comes from a plasmid or a chromosome. The random forest classifier algorithm was trained with the *RandomForestClassifier* function from *scikitlearn* library (16) in Python 3.6, using the seven features described above. The number of random decision trees was kept to 500, as out-of-bag error estimate (*i.e*., the internal error of individual decision trees during the training process) did not significantly decrease when using more trees. The global classification method of PlasForest is described in Fig. 3.

**Figure 3:**
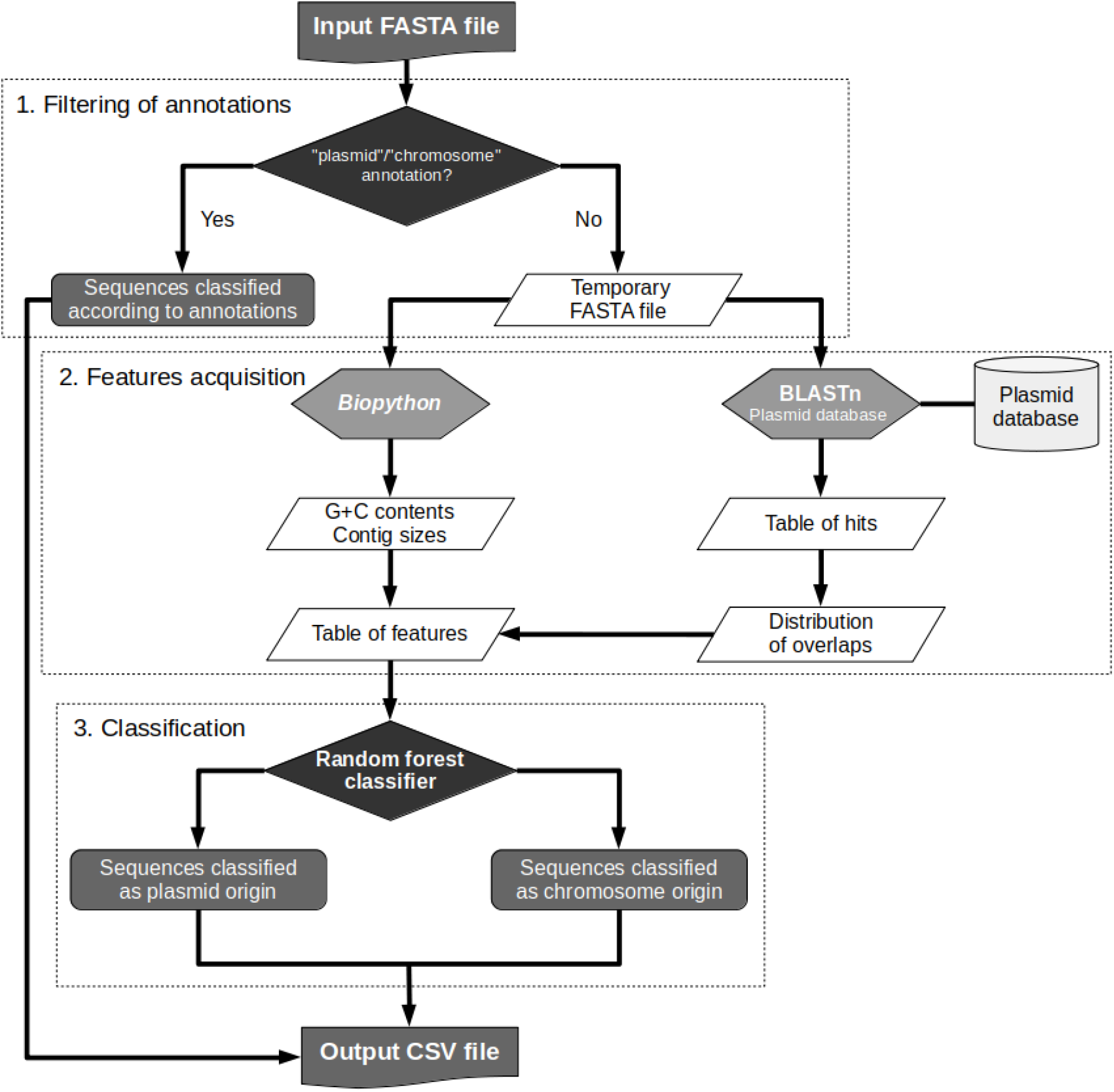
General method of classification implemented in PlasForest.

#### 2.2.3 Sensitivity of the classifier

We tested the sensitivity of PlasForest (i) to the composition of the plasmid database and (ii) to the composition of the balanced training set, by performing two independent bootstrap analyses. To assess the importance of the composition of the plasmid database, we resampled the plasmid database with 50 different seeds. We then computed new features for each contig of the balanced training and testing set. Classifiers were trained on the resulting balanced training sets, and their performances were measured on the resulting balanced testing sets. To test the sensitivity to the composition of the balanced training set, we resampled the balanced training set 50 times, while the balanced testing set did not change. We trained classifiers on the resulting balanced training sets, and measured their performances on the balanced testing set.

### 2.3 Measure of classification performances

#### 2.3.1 Indices of binary classification performance

In order to measure the performance of our trained algorithm to correctly identify plasmid sequences and to compare its performance to other available tools, we computed indices derived from the confusion matrix that are commonly used in binary classifications. Sensitivity (sometimes indicated as recall) is the fraction of positive data (in our case, plasmid contigs) which has been correctly identified as positive and allows to measure the false negative error rate. Precision (also indicated as the positive predictive value) corresponds to the fraction of positive results that are actually true positives and allows to measure the false positive error rate. A good classifier should be able to optimize both sensitivity and precision *i.e*. in our case, identify as many plasmid contigs as possible without misidentifying chromosome contigs as plasmids. For this reason we calculated “composite” indices that reflect the overall performance of the classifier. F1 score corresponds to the harmonic mean of sensitivity and precision: it therefore ranges from 0 (*i.e*., either precision or sensitivity or both are null) to 1 (*i.e*., there are neither false positives nor false negatives). Accuracy corresponds to the fraction of correct predictions, *i.e*. the fraction of true positives and true negatives (in our case, plasmid contigs identified as plasmids and chromosome contigs identified as chromosomes). F1 score does not take into account negative predictions and accuracy can be biased when one of the two classes is rare compared to the other. We also calculated *Matthews Correlation Coefficient* (MCC). This metric corresponds to a correlation coefficient between the observed and the predicted classifications and is generally regarded as a balanced measure that can even be used if classes are of very different sizes (17). Values range between +1 for a perfect prediction, 0 for a random prediction, and −1 for a prediction in total disagreement with the observed data.

#### 2.3.2 Comparison with other softwares

PlasForest, PlasFlow (8) and cBar (9) were run on the full balanced testing set (157,364 contigs, from 50pb to 3.7Mb). The classification outputs were compared to the annotations attributed in RefSeq to compute performance indices for both global comparison and comparison by contig size categories.

##### 2.3.2.1 Prediction of plasmid sequences using PlasmidFinder

To identify plasmid sequences using PlasmidFinder, we uploaded the sequences of the balanced testing set to the PlasmidFinder webserver (https://cge.cbs.dtu.dk/services/PlasmidFinder/) (6). As PlasmidFinder is specialized on Enterobacteriaceae and Gram-positive bacteria plasmids, it was run on a subset of this dataset containing only these taxa.

##### 2.3.2.2 Prediction of plasmid sequences using cBar

cBar 1.2 was downloaded from http://csbl.bmb.uga.edu/~ffzhou/cBar/ and the program was run on the complete balanced testing set (9).

##### 2.3.2.3 Prediction of plasmid sequences using PlasFlow

The version 1.1 of PlasFlow was downloaded from https://github.com/smaegol/PlasFlow and the program was run on the complete balanced testing set (8). Taxonomic assignments were not taken into account to assess the performance of the classification. When the algorithm assigned sequences as “unclassified”, they were not considered as positive or negative results and were not used to assess the performance of the classification.

### 2.4 Processing time

We measured the processing time of PlasForest, and we compared it with PlasFlow, on all the representative genomes at the assembly level ‘Contig’ published between July 1st, 2020 and July 31st, 2020 on NCBI Refseq Genomes FTP. PlasForest and PlasFlow were run on these 108 genomes, representing 1897 contigs, on a computer with 6 cores and 16GB of RAM. The whole identification process was completed in 18 minutes for PlasForest and in 8.5 minutes for PlasFlow. PlasForest and PlasFlow agreed on the classification of 970 contigs (957 classified as chromosome, 13 as plasmid), and disagreed on the classification of 515 contigs (the rest being labeled “unclassified” by PlasFlow).

## 3. Results

PlasForest is a tool that allows to assign contigs in genomic datasets to plasmid or chromosome by using a random forest classifier on variables extracted from a homology search. We tested the sensitivity of its performances to various changes in the training process, and we compared its performances with other classical plasmid identification softwares on large genomic datasets.

### 3.1 Pipeline description

A pipeline executing all analyses required for PlasForest was encoded in Python 3.6. This pipeline performs several steps described in Figure 3. First, it filters query contigs by their annotations, to avoid re-identification of sequences already described as chromosome or plasmid (step 1). If the user wants to re-assign contigs with Plasforest, this step can be skipped with option *-r*. Feature acquisition (step 2) consists of submitting the filtered sequences to BLASTn against a local copy of the plasmid database, and calculation of the overlaps between query and subject sequences. Features are then computed for each query contig on its sequence and the distribution of overlaps. The classification (step 3) passes features to the random forest classifier, which outputs the predicted identification for each query contig. The output can also include the best hits that were found in the plasmid database with option *-b* and/or the features used for classification with option *-f*. Progression of the pipeline can be displayed if run with option *-v*.

### 3.2 Reliability of the classification method

We aimed to test the reliability of PlasForest by measuring its sensitivity to qualitative changes. We tested whether adding or removing features in the training of the classifier would yield better performances. We also tested whether the classifier could maintain high performances when resampling the training dataset and the plasmid database.

The classifier was trained several times on the balanced training set, by feeding it with different combinations of features. Predictions of the classifiers were made on the balanced testing set. The classification based on all seven features (*average overlap, contig size, G+C content, max overlap, median overlap, number of hits, variance overlap*) showed highest MCC, and the three variables that showed the highest feature importance are the *maximal overlap*, the *number of hits*, and *contig size* (Fig. 2C).

The distribution of performances of PlasForest obtained after each bootstrap procedure is displayed in Fig. 4. As expected, the variation of performance of PlasForest is low for the 50 iterations of the bootstrapped data. The performance of PlasForest presents a very low sensitivity to the resampling of the training set (see Fig. 4A). However, the performance of PlasForest decreases substantially when the composition of the plasmid database is resampled (see Fig. 4B). Removing sequences from the plasmid database can therefore significantly diminish the ability of PlasForest to correctly assign contigs.

**Figure 4:**
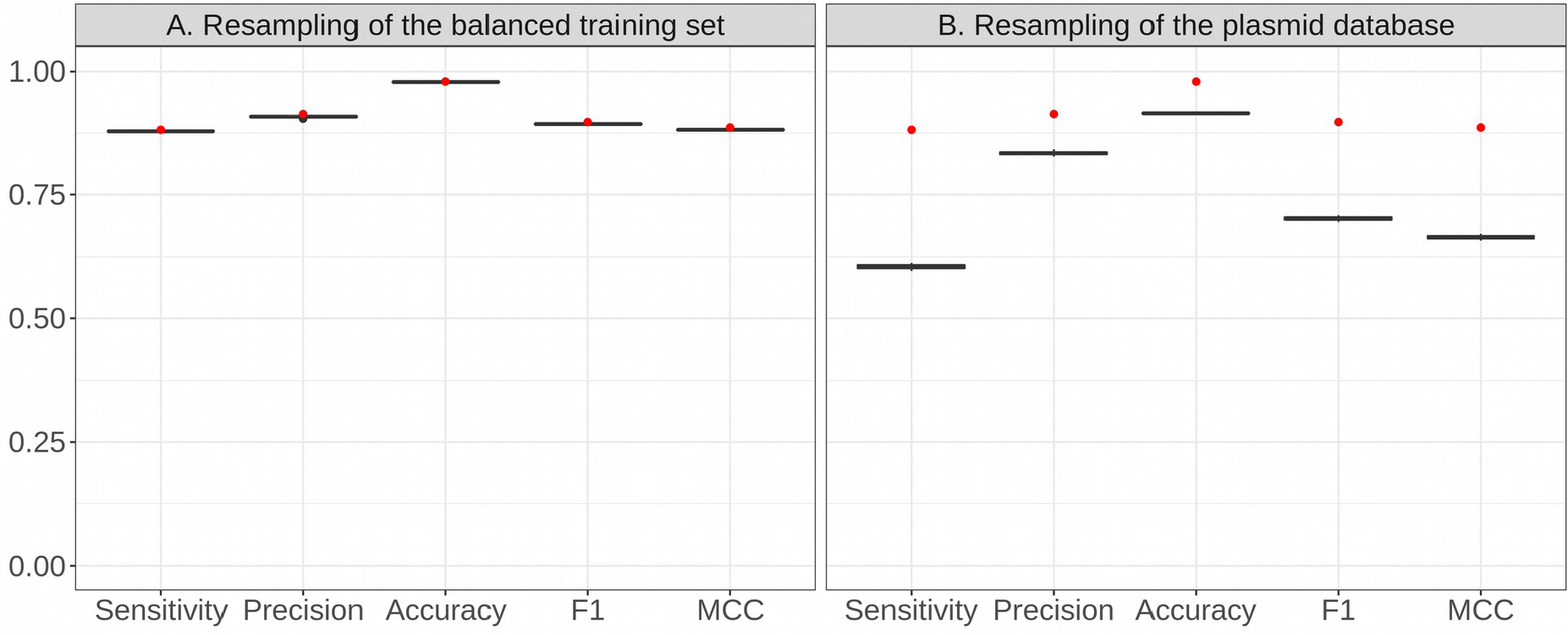
Sensitivity of PlasForest to resampling of the balanced training set and the plasmid database. The initial performances of PlasForest on the balanced testing set are displayed with red dots. The distribution of performances for PlasForest when resampling 50 times either the plasmid database or the balanced training set are displayed in grey boxes.

### 3.3 Adaptability of the performances on the external validation dataset

The dataset on which the classifier was trained and validated might be biased (assembled genomes) and does not represent the taxonomic diversity of bacteria. We then verified the adaptability of PlasForest to taxa on which it had not been trained. We ran PlasForest on the external validation dataset which was drawn from randomly cut genomes that were left apart from the holdout validation process. Overall, PlasForest made less correct predictions on the external validation dataset than it did on the balanced testing set, with an overall sensitivity of 72.1% and an overall precision of 87.7%. Yet, PlasForest keeps a good performance, with a MCC=0.781 and an accuracy of 97.2%. PlasForest thus does not overfit the data.

### 3.4 Benchmark of plasmid identification methods

We compared the predictions of PlasForest on our balanced testing dataset (see Fig. 5), with those of two classical k-mer-based classification methods (cBar (9) and PlasFlow (8)) and one classical homology-based classification method (PlasmidFinder (6)). We chose not to include PlasmidSeeker (10) to our analysis, because it requires to know the species from which the genome comes and therefore cannot be used on a wide scale.

**Figure 5:**
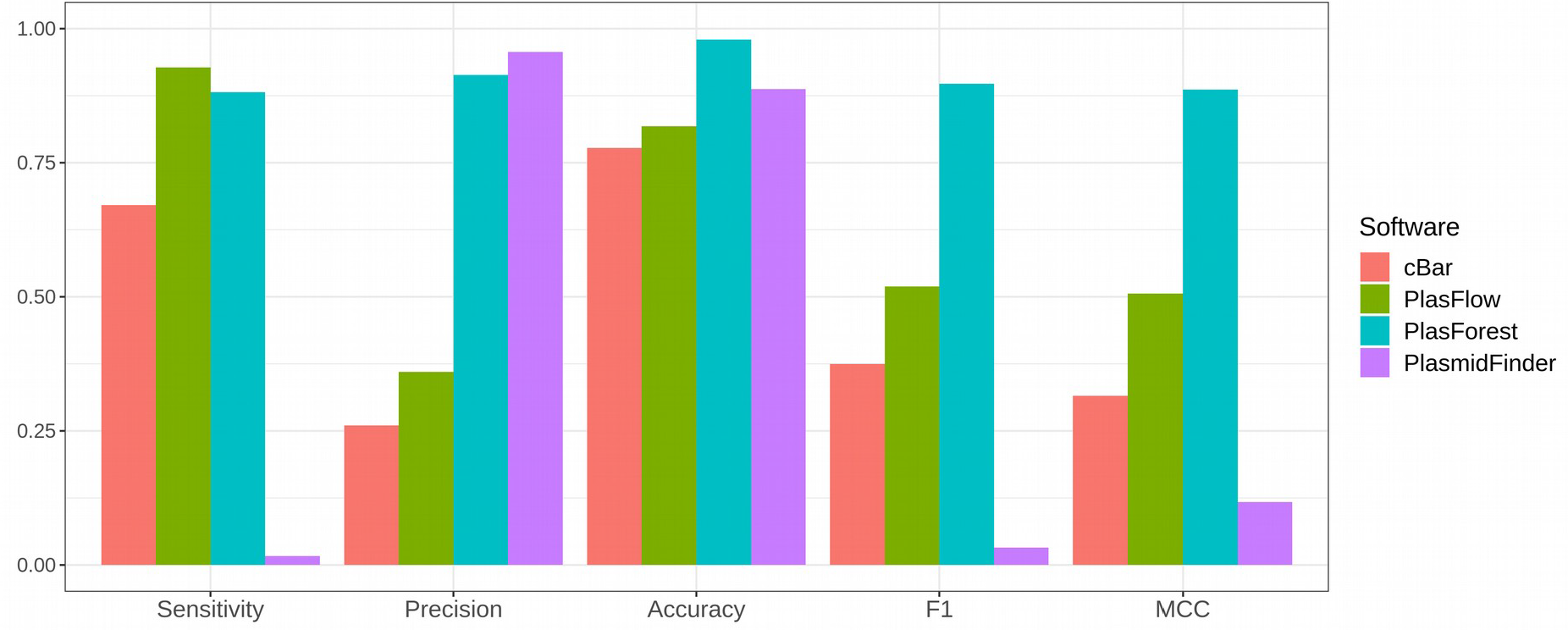
General performances of all tested softwares against the testing set. Bars represent the performances of the four tested softwares against the balanced testing set: cBar is displayed in red, PlasFlow in green, PlasForest in blue, and PlasmidFinder in purple. Indices of performance are scores of binary classification, based on the confusion matrix.

PlasForest is overall the second most sensitive classifier, being able to predict 88.2% of plasmid contigs whereas PlasFlow can predict 92.8% of plasmid contigs. However, for sequences above 50kb, PlasForest is as sensitive as or more sensitive than PlasFlow with 95.9% of plasmids correctly predicted. PlasmidFinder had the highest precision for predictions on the *Enterobacteriaceae* and Grampositive bacteria dataset, followed by PlasForest (with respectively 4.5% and 8.6% of sequences incorrectly predicted as plasmids). For contigs above 50kb, the precision of PlasForest is comparable with that of PlasmidFinder (3.3% of false positives). On these two indices, PlasForest is therefore globally the second best software, and even the best above 50kb. On the contrary, PlasFlow suffers from low precision, especially below 50kb (up to 79% of false positives), and PlasmidFinder suffers from low sensitivity for all contig sizes (down to 2.9% of true positives). PlasForest therefore outperforms all other softwares on the composite indices, with the highest MCC (globally 0.886, and even 0.959 for contigs above 50kb), the highest F1 score (globally 0.897, and even 0.962 for contigs above 50kb), and the highest accuracy (globally 98.0%, and even 99.3% of correct attributions for contigs above 50kb) for all contig sizes. Even for very short contigs (under 1kb) for which k-mers-based methods usually have poor results, PlasForest remains a reliable classifier, with MCC=0.775 and an accuracy of 96.0% (respectively MCC=0.146 and 35.3% for PlasFlow).

## 4. Discussion

PlasForest is a homology-based approach, combined with machine learning, to detect plasmids in contig and scaffold genomes. Its operating principle is to seek homologies between query contigs and a large plasmid database, and then to assign a plasmid/chromosome identity to queries with a random forest classifier.

Here we showed that PlasForest is able to deal with large datasets without prior knowledge of the taxonomic background, and that it identifies plasmids with both a high sensitivity and a high precision in unassembled genomes. All the plasmid identification software tested had either a low false negative error rate or a low false positive error rate. PlasForest did not have the lowest individual error rates, but it optimized the tradeoff between sensitivity and precision. In terms of biological material, PlasForest should be able to deal with contig and scaffold genomes as well as assembled genomes. For contig and scaffold genomes, PlasForest has been trained and tested on artificial draft genomes and has good performance on them (overall accuracy of 98.0% and MCC=0.886; and up to an accuracy of 99.3% with MCC=0.959 on contigs above 50kb). Table 1 summarizes the four plasmid identification methods that we compared in this article.

**Table 1:** Summary of the four plasmid identification methods/tools tested in this study.

| Software | PlasmidFinder | cBar | PlasFlow | PlasForest |
| --- | --- | --- | --- | --- |
| Machine learning algorithm | - | + | + | + |
| K-mers-based | - | + | + | - |
| Homology-based | +++ | - | - | ++ |
| Applicable to many taxa | - | + | + | + |
| Applicable to metagenomic datasets | - | ++ | +++ | - |
| Sensitivity | - | ++ | +++ | ++ |
| Precision | +++ | + | + | ++ |

K-mer-based approaches that are able to identify specific genomic signatures of chromosomes and plasmids were traditionally the most reliable plasmid identification methods. The number of possible k-mers increases exponentially with the size of k-mers. Therefore, accurately estimating the frequency of all possible k-mers of a given size, even for short k-mers, requires large contigs. For example, estimating the frequency of all 256 possible 4-mers in a contig should require thousands of bases, while the size of the shortest known plasmids is below 1kb (18). Thus, PlasForest outperforms those methods, especially on short contigs, because the quality of the genomic signature is much less dependent on contig size with homology-based features than with k-mers. Some k-mer-based approaches compensate this limitation, but can only be used in a very restricted context: for example, PlasmidSeeker (10) achieves up to 100% sensitivity and 99.8% specificity in whole genome sequencing reads by ruling out as chromosomal the k-mers which are shared with a complete, reference assembly. However, the use of PlasmidSeeker on a wide scale is practically infeasible: it requires to know the species from which the genome comes from and that at least one genome has been assembled for this species (10).

The high precision of PlasForest is inherited from its homology-seeking basis. Though homology-based approaches generally show little sensitivity, due to their inability to detect unrelated plasmids, the performances of PlasForest are significantly higher than those of the other homology-based method tested (PlasmidFinder (6)). One of the reasons why PlasForest is as sensitive as (or even more than) k-mer-based approaches is that it aggregates measures of homologies in a classifier. Thus, it not only considers the presence of homologies, but it also measures the quality and diversity of these homologies and this improves the accuracy of the identification process.

Many bacterial species remain uncultivable but an increasing number of metagenomics tools have allowed access to genomic data in environmental samples without the necessity of obtaining pure cultures. Other identification tools specially designed for such data (*e.g*., PlasFlow (8)) offer the opportunity of taxonomic assignment of the sequence identified as belonging to plasmids. During the development of PlasForest, we addressed the classification of contigs from genomic (and not metagenomic) data into plasmids or chromosomes. The pipeline of PlasForest offers the possibility to identify which plasmids from the database have the strongest homology with the query. Though, it should be noted that the taxonomic assignment can strongly depend on (i) the plasmid host range and (ii) horizontal gene transfer events that are yet massively undetermined in bacteria. Any attempt (from PlasForest or from any other method) of taxonomic assignment for broad host range plasmids is thus at least imprecise and can sometimes be impossible (19).

Finally, in its current state, PlasForest cannot be applied to detect plasmids in metagenomes. Sampling metagenomic data generates short reads (current standards at 2×150bp paired-end, see (12)), for which no current identification method is able to produce accurate assignment (here for PlasForest, ca. 60% sensitivity for contigs of 1kb or less). In theory, this low performance could be overcome by a partial assemblage of raw reads. However, as PlasForest mostly relies on homologies with a plasmid database, the vast majority of plasmids it is able to detect are related to those of its database. Unrelated plasmids with narrow host ranges, especially if the host is uncultivated, cannot be detected through this method so far. In order to achieve homology-based identification of plasmids in metagenomes, new datasets are required. Some sequences from assembled plasmidomes (some being already available on MG-RAST database (20) could be incorporated in the plasmid database. But most importantly, when plasmids can be mechanically separated from chromosomes in metagenomic samples (*e.g* (21)), both chromosomes and plasmids should be sequenced: PlasForest could then be trained on annotated metagenomic datasets. As these datasets become more and more available, PlasForest should become able to detect plasmids in metagenomes.

In its current state, among all softwares tested, PlasForest is the best identification method for plasmids in contigs and scaffolds. We released PlasForest as a user-friendly pipeline, including the trained classifier and the plasmid database, directly available on GitHub (https://github.com/leaemi-liepradier/PlasForest). Further releases (at least on a half-year basis) will include a plasmid database updated with the new plasmid sequences submitted to public repositories, and a new classifier trained on this database. This complemented database should be regularly trimmed, in order to keep it of reasonable size without decreasing the performance of PlasForest. Especially in order to identify plasmids in metagenomic data with the same accuracy as k-mer-based approaches, plasmidomes should be included in the plasmid database and the classifier should also be trained on annotated metagenomic data.

## 5. Availability

PlasForest was released on GitHub (https://github.com/leaemiliepradier/PlasForest) and is available under GPLv3 license.

## Acknowledgments

We thank Martijn Callens and Emira Cherif for his comments on a previous version of the manuscript.

